# Establishment in Culture of Expanded Potential Stem Cells

**DOI:** 10.1101/124479

**Authors:** Jian Yang, David J. Ryan, Wei Wang, Jason Cheuk-Ho Tsang, Guocheng Lan, Hideki Masaki, Xuefei Gao, Liliana Antunes, Yong Yu, Zhexin Zhu, Juexuan Wang, Aleksandra A. Kolodziejczyk, Lia S. Campos, Cui Wang, Fengtang Yang, Zhen Zhong, Beiyuan Fu, Melanie Eckersley-Maslin, Michael Woods, Yosuke Tanaka, Adam C. Wilkinson, James Bussell, Jacqui White, Ramiro Ramirez-Solis, Wolf Reik, Berthold Göttgens, Sarah A. Teichmann, Hiromitsu Nakauchi, Xiangang Zou, Liming Lu, Pentao Liu

## Abstract

Mouse embryonic stem cells are derived from *in vitro* explantation of blastocyst epiblasts^1,2^ and contribute to both the somatic lineage and germline when returned to the blastocyst^3^ but are normally excluded from the trophoblast lineage and primitive endoderm^4–6^. Here, we report that cultures of expanded potential stem cells (EPSCs) can be established from individual blastomeres, by direct conversion of mouse embryonic stem cells (ESCs) and by genetically reprogramming somatic cells. Remarkably, a single EPSC contributes to the embryo proper and placenta trophoblasts in chimeras. Critically, culturing EPSCs in a trophoblast stem cell (TSC) culture condition permits direct establishment of TSC lines without genetic modification. Molecular analyses including single cell RNA-seq reveal that EPSCs share cardinal pluripotency features with ESCs but have an enriched blastomere transcriptomic signature and a dynamic DNA methylome. These proof-of-concept results open up the possibility of establishing cultures of similar stem cells in other mammalian species.

---

We sought to establish cultures of new stem cells from cleavage stage mouse embryos. Under such a culture condition, we speculated that the self renewing stem cell population might have expanded potential as the cells of 4-cell (4C) or 8-cell (8C) embryos or the individual blastomeres retain the potential to differentiate to both the trophoectoderm (TE) and the inner cell mass (ICM)^7–10^. In order to prevent blastomeres from further differentiation and to derive stem cell lines from these cells, we speculated that modulation of the key signaling pathways implicated in the earliest stages of embryonic development might be a rate-limiting step. Genetic and developmental studies have revealed the key roles of conserved mitogen-activated protein kinases (MAPKs), Src, and Hippo pathways in the segregation of the TE and ICM lineages; and furthermore, how disrupting them causes developmental arrest^11–17^. In addition, Wnt signaling is known to be a key orchestrator of the earliest development stages of vertebrates^18^, and to be involved in the development of mouse preimplantation embryos and trophoblasts^19–22^. Recent advances have uncovered important functional interactions between Wnt and MAPK pathways via Yap, the key Hippo pathway downstream effector^23–25^. We therefore selected inhibitors to simultaneously target these pathways or kinases to block development for the derivation of novel stem cell lines.

In order to target MAPKs, we used Mek1 inhibitor PD0325901, JNK Inhibitor VIII (for Jun N-Terminal Kinase) and p38 inhibitor SB203580. We chose A-419259, a potent pyrrolopyrimidine inhibitor, to block activities of Src family kinases^26^. To modulate Wnt signaling, XAV939 was used to stabilize AXIN1, the concentration-limiting component of the β-catenin and Yap destruction complex^27,28^. XAV939 may also suppress Yap activities via angiomotin^25^, and has been shown to improve culturing pluripotent stem cells^29,30^. Finally, we included a GSK3 inhibitor, CHIR99021, and leukemia inhibitory factor (LIF). Although, the pro-pluripotency role of GSK3 inhibition (higher Wnt activities) appears to be partially redundant when PD0325901 and LIF are present^31–34^, CHIR99021 improves metabolic and biosynthetic processes and therefore cell culture robustness^35^. LIF may promote the rare totipotent cells in mouse ESC culture^36^. We hereafter named the medium that contains the aforementioned inhibitors and LIF as EPSC Medium or EPSCM, which was used in subsequent experiments, unless otherwise stated.

We first investigated whether we could derive cell lines directly from single blastomeres of an 8C embryo in EPSCM, as previous attempts to culture mouse blastomeres only produced standard ESC lines^37^. We seeded each blastomere into a well of a 96-well plate with feeders (Fig. 1a). In the following days, in 9 out of 32 wells with EPSCM, blastomeres proliferated, and by day 12, formed large outgrowths consisting predominantly of Oct4^+^/Cdx2^−^ cells (Fig. 1a). Stable EPSC lines could be established in EPSCM from most of the outgrowths (Fig. 1a). By contrast, no outgrowths were obtained from blastomeres in either M15 (15% serum plus LIF, or mouse ESC medium) (n=32) or 2i/LIF (n=32) on feeders, or in feeder-free conditions in any medium (n=96).

**Figure 1.**
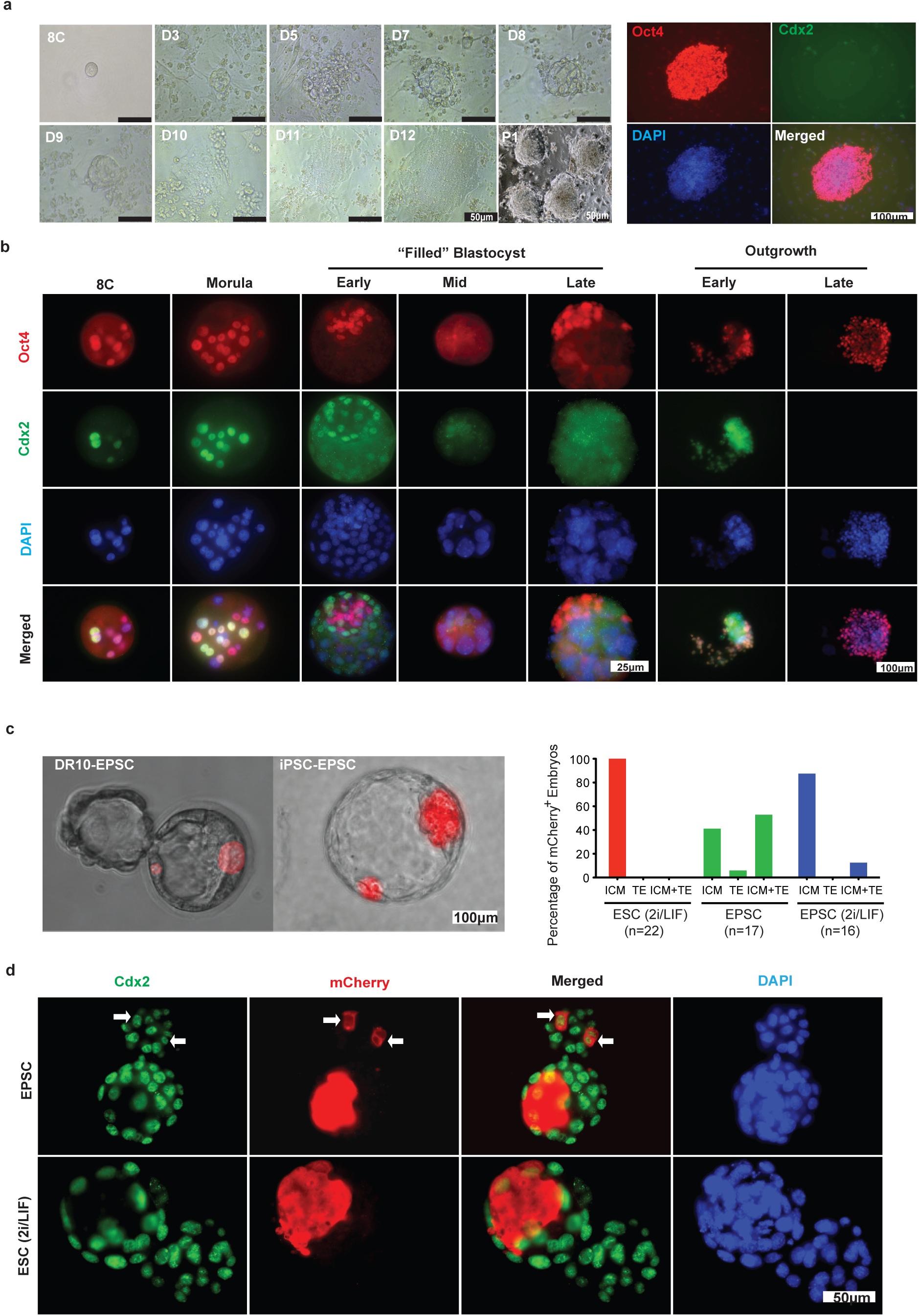
Derivation of EPSC lines from single blastomeres or preimplantation embryos. **a**. Derivation of EPSC lines from single blastomeres. Individual 8C blastomeres were seeded in each well of a 96-well plate on feeders in EPSCM. Temporal progression of the blastomere was imaged. P1: passage 1 of a primary outgrowth. Right panels: immunostaining for Oct4 or Cdx2 in cells of a primary outgrowth from a single blastomere. Most cells were Oct4^+^ and Cdx2^−^. **b**. Progression of an 8C embryo in EPSCM. Oct4 and Cdx2 expression was detetced by immunofluorescence. Early, mid or late “filled” blastocysts refer to those in EPSCM for 72-96 hours, 5-6 or 7-8 days, respectively. In the mid-filled-blastocysts, most if not all cells do not express either nuclear Oct4 or Cdx2. Oct4^+^ cells eventually emerged from the edge of the late “filled” blastocysts and form primary outgrowths. **c**. TE contribution in the blastocyst of EPSCs (mCherry^+^) derived from preimplantation embryos or converted from iPSCs (merged live images of phase and mCherry fluorescence). The chart on the right describes the percentages of TE contribution of cells in EPSCM or 2i/LIF. **d**. The blastocysts (hatching) developed from morulas injected with mCherry^+^ ESCs or EPSCs were co-stained for mCherry (in the cytoplasm) and Cdx2 (in the nucleus). Embryos were stained with DAPI for the nucleus. The arrows indicate cells expressing both Cdx2 and mCherry. The experiments were repeated at least three times.

Next, we examined whether a cleavage stage embryo could progress developmentally in EPSCM. Surprisingly, these 4C or 8C embryos in EPSCM initially appeared to progress to blastocyst-like structures (Extended Data Fig. 1a, b). However, in EPSCM blastocysts, the blastocoel was obliterated by day 6 and was filled with cells, generating a structure reminiscent of an enlarged morula, or a “filled” blastocyst (Extended Data Fig. 1a, b). The “filled” blastocysts eventually attached and hatched, and after day 10, primary outgrowths appeared (Extended Data Fig. 1a). EPSC lines were established from the primary outgrowths in EPSCM with an efficiency of approximately 20% under feeder-free condition, and up to 100% on SNL feeder cells. We investigated Oct4 and Cdx2 expression in the developing embryos in EPSCM (Fig. 1b) and in the control medium M15 (Extended Data Fig. 1c). Immunostaining revealed a gradual loss of Oct4 and Cdx2 expression in the mid-“filled” blastocysts, which eventually led to a temporary “blank” state, where most cells had large nuclei, but almost none expressed nuclear Oct4 or Cdx2 (Fig. 1b and Extended Data Fig. 1d, e). This transient “blank” state appears to bear certain similarities with some of the early blastomeres (4C embryo in M15 in Extended Data Fig. 1c), or with the 2C-like ESCs where Oct4 protein expression is absent^38^. Genetically, Oct4 expression is not essential for the establishment of totipotency^39^. Further analysis of the “filled” blastocysts indicated that the cells appeared to be undergoing a complex reprogramming process involving cell proliferation and apoptosis, as many were positive for Ki67 (Extended Data Fig. 1d) and cleaved Caspase 3 (Extended Data Fig. 1e). This observation, and particularly the properties and potential of the individual cells within the “filled” blastocysts, warrant further investigation. Notably, in the primary outgrowths, some cells initially expressed both Oct4 and Cdx2 (Fig. 1b), as was the case with the co-expression of the two transcription factors in many 4C-8C blastomeres (Fig. 1b and Extended Data Fig. 1c). Eventually the outgrowth cells and the established EPSC lines expressed only Oct4 (Fig. 1b). In contrast, in 2i/LIF, only 5 out of 54 embryos (8C) reached the blastocyst stage and all died soon after.

The fact that embryo cells in EPSCM went through transient loss of Oct4 and Cdx2 expression, and also that some cells in the primary outgrowths initially expressed both Oct4 and Cdx2, implied that EPSCs might retain some blastomere features, even though they were morphologically similar to standard ESCs. We characterised two cells lines (DR10-EPSCs and DR25-EPSCs) derived *de novo* in EPSCM from 8C embryos. These cells, which expressed pluripotency genes at levels comparable to standard ESCs, had a normal karyotype, were able to form mature teratomas, and contributed to both the somatic lineage and the germline in chimeras (Extended Data Fig. 1f, g, h, i, j, k). Remarkably, once injected into morulas, both EPSCs (mCherry^+^) contributed to the ICM and the TE in the blastocysts (Fig. 1c and Extended Data Fig. 1l).

We subsequently cultured standard ESCs (AB2.2 and E14Tg2a (ref. 40,41)) or induced pluripotent stem cells (iPSCs) reprogrammed from MEFs in EPSCM, which were previously maintained in M15 or 2i/LIF. After five passages in EPSCM, these ESCs or iPSCs appeared to acquire the expanded potential to contribute to the TE once injected into recipient embryos (Fig. 1c and Extended Data Fig. 1l). Importantly, the donor cells from injected EPSCs expressed Cdx2 (Fig. 1d). Notably, once EPSCs were returned to 2i/LIF for as few as three passages, a much lower contribution to the TE was observed (Fig. 1c), indicating that EPSCM is necessary to sustain the expanded potential, and that EPSCs could re-acquire naïve ESC characteristics in an ESC culture condition. We named ESCs or iPSCs consecutively cultured in EPSCM for five passages as ESC-EPSCs or iPSC-EPSCs. The ESC-EPSCs could be cultured for over 30 passages, and retained robust levels of reporter GFP expression from the *Oct4* and *Rex1* loci^42,43^ (Extended Data Fig. 1m, n). They expressed comparable levels of mRNAs of pluripotency and lineage-specific genes as in ESCs (Extended Data Fig. 1o); they preferentially used the *Oct4* distal enhancer in lieu of the proximal enhancer^44,45^ (Extended Data Fig. 1p); they differentiated to all three somatic germ layers *in vitro* (Extended Data Fig. 1q) and to the germline in chimeras (Extended Data Fig. 1r), and also showed efficient gene targeting (around 50%) at the *Rosa26* locus^46^. Epigenetically, female EPSCs had two active X chromosomes (Extended Data Fig. 1s). EPSCs thus shared the cardinal features associated with pluripotency. Biochemically, targeting the pathways and kinases known to be important in the earliest developmental stages by the inhibitors in EPSCM resulted in the effective modulation of their activities (Extended Data Fig. 1t). Notably, EPSCs had increased Axin by XAV939 (Extended Data Fig. 1t), which led to higher levels of phosphorylated-β-catenin and lower active β-catenin in the nucleus (Extended Data Fig. 1u). Consequently, EPSCs did not have detectable Wnt activity in Topflash assay (Extended Data Fig. 1v), even though EPSCM contained the GSK3 inhibitor CHIR99021, demonstrating XAV as a potent Wnt inhibitor. EPSCs were responsive to LIF or Jak/Stat signaling (Extended Data Fig. 1w) but were resistant to FGFRi and ALK5i, similar to ESCs in 2i/LIF (Extended Data Fig. 1x).

We further determined the *in vivo* differentiation potency of EPSCs by implanting injected recipient 8C embryos after they had developed to the morula (at CRI) or blastocyst (at the Sanger) stage. The embryos were subsequently collected around 6.5dpc for assessment of contribution. In about half of the chimeras (59/113), donor mCherry^+^ cells were found in the extraembryonic ectoderm region that was stained positively for Elf5 (Ref. 47) (Fig. 2a, and Extended Data Fig. 2a). In contrast, 2i/LIF ESCs rarely contributed to this region. To ensure reproducibility of EPSC contribution in chimeras, we independently converted another GFP-expressing ESC line in EPSCM at the University of Tokyo. These GFP^+^ EPSCs, but not the GFP^+^ 2i/LIF ESCs, once injected into recipient blastocysts, contributed to the extraembryonic ectoderm region in 6.5dpc chimeras (11/68) (Fig. 2a. Top panel, live images).

**Figure 2.**
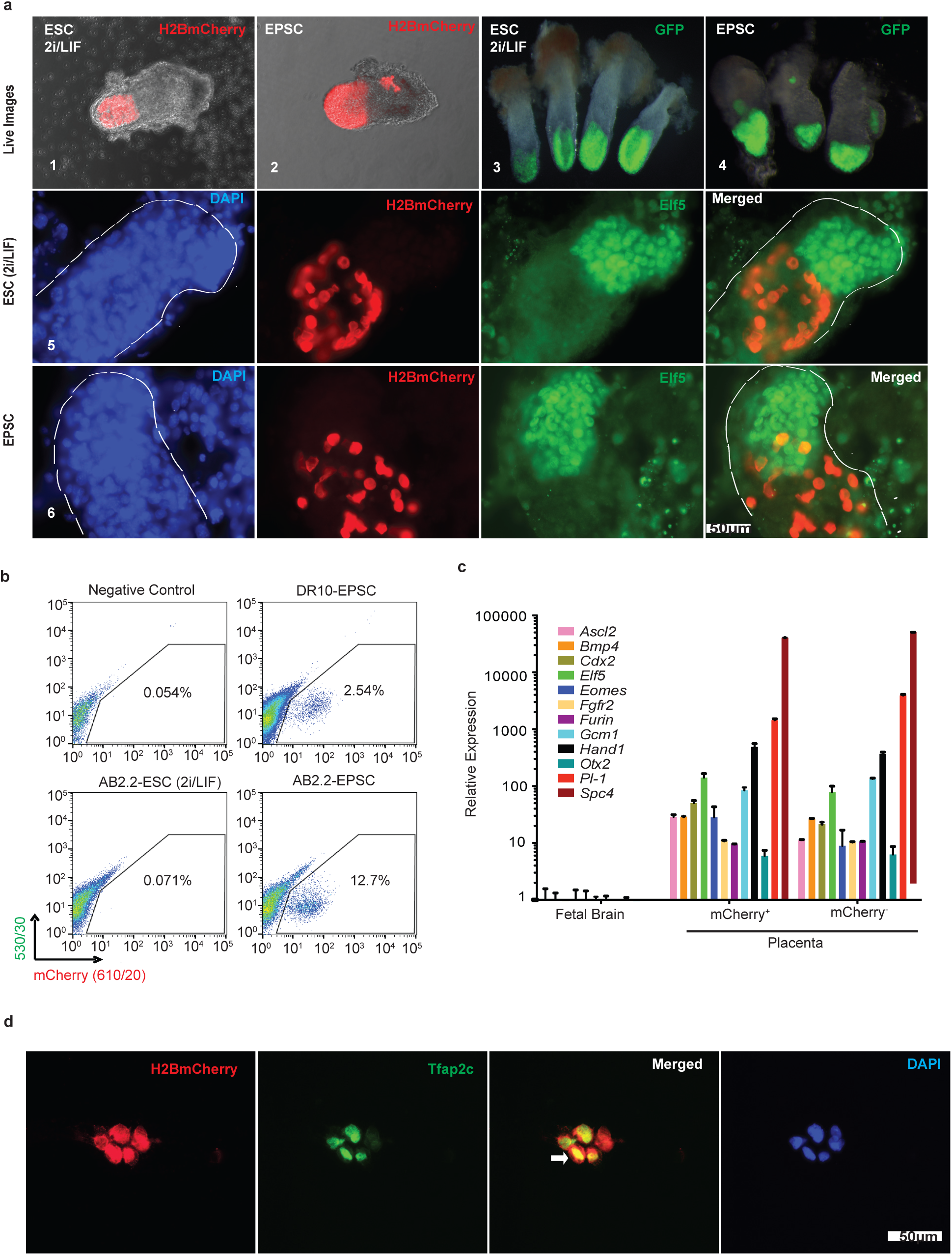
Contribution of EPSCs and ESCs to trophoblasts in chimera embryos. **a**. Whole-mount fluorescence imaging of representative 5.5-6.5dpc ESC or EPSC chimeric embryos produced in Cambridge, UK (AB2.2-H2BmCherry), or at University of Tokyo (BDF1-EGFP#4). The top panels include four live images. The rest (embryos 5 and 6) are immunofluorescence staining of chimeras for mCherry and Elf5. DAPI stains the nucleus. Embryos are marked by thin dashed lines. **b**. Detection of donor mCherry^+^ placenta cells from EPSC 14.5dpc chimeras. The contribution to the placenta from 2i/LIF ESCs was low. We used the GFP channel to exclude autofluorescence. **c**. Expression of trophoblast genes in sorted donor mCherry^+^ and host mCherry^−^ placenta cells from an EPSC 14.5dpc chimera. Expression was normalized to fetal brain expression. Data are the mean ± s.d. **d**. Detection of Tfap2c in sorted mCherry^+^ cells from 14.5dpc EPSC chimera placenta. The sorted cells were cytospun to polylysine coated slides, and stained for mCherry and Tfap2c. The arrow points to mCherry^+^ cells that are Tfap2c^+^. Blood cells were gated out. The experiments were repeated at least three times.

In order to examine the contribution of EPSCs in the placenta, we analysed 14.5dpc chimeras. Whole-mount fluorescence examination of the chimeras indicated mCherry^+^ donor cell contribution to the embryo proper and possibly the extraembryonic tissues (33 of 71 embryos for DR10-EPSCs, and 23 of 41 for DR25-EPSCs, 21 out of 38 for AB2.2-EPSCs and 16 of 28 for AB2.2-2i/LIF ESCs) (Extended Data Fig. 2b). Flow cytometry analysis of dissociated single cells from the placenta confirmed the presence of descendants of mCherry^+^ EPSCs (Fig. 2b). We sorted mCherry^+^ (donor) and mCherry^−^ (host) placenta cells for gene expression, which revealed that both groups of cells expressed comparable levels of trophoblast genes, such as *Ascl2* (*Mash2*), *Gcm1, Pl-1*, and *Hand1* (ref. 48) (Fig. 2c). By contrast, flow cytometry failed to detect a substantial number of descendants of AB2.2-2i/LIF ESCs in the placenta (Fig. 2b).

Many mCherry^+^ placenta cells from EPSC chimeras were polyploid in DNA content analysis (Extended Data Fig. 2c), similar to mCherry^−^ cells, as trophoblasts could be detected as 2N, 4N, 8N and so on, due to endoreplication and cell fusion in placenta development^49^. At the cellular level, immunostaining of placenta sections identified mCherry^+^ cells in EPSC chimeras, but not in the placenta of 2i/LIF ESCs (Extended Data Fig. 2d). Notably, mCherry^+^ placenta cells were stained positively for transcription factor Tfap2c (Extended Data Fig. 2d), which is expressed in spongiotrophoblasts and other types of trophoblasts^50^, and is one of the factors for the generation of induced trophoblast stem cells^51,52^. We further FACS-purified mCherry^+^ placenta cells from the EPSC-chimeras, cytospun and immunostained them for additional trophoblast markers. Besides Tfap2c, trophoblast markers Gcm1, Ezrin and Cytokeratin 7 (CK7) could clearly be detected in the sorted mCherry^+^ placenta cells (Fig. 2d, and Extended Data Fig. 2e).

We wished to determine and exclude the possibility that mCherry^+^ placenta cells had gone through cell fusion between the donor EPSCs and host trophoblasts at any stage of development. Therefore, we injected H2B-mCherry^+^ AB2.2 EPSCs into host embryos that were genetically marked (The *SleepingBeauty Transposase-SB10* was targeted to the *Rosa26* locus)^46^. Genotyping the sorted mCherry^+^ placenta cells (14.5dpc) for the presence of *mCherry* or *SB10* genomic DNA confirmed robust *mCherry* DNA amplification, but no *SB10*, whereas in the mCherry^−^ cells *SB10* DNA could be readily detected (Extended Data Fig. 2f), thus genetically excluding cell fusion events that could have produced mCherry^+^ placenta cells. In the mCherry^−^ placenta cells, minor *mCherry* DNA amplification was noticed (Extended Data Fig. 2f), indicating a small number of mCherry^+^ donor cells becoming mCherry^−^, due to the silencing of the CAG promoter that controlled mCherry expression.

The yolk sac of 14.5dpc chimeras consists of cells derived from the extraembryonic mesoderm, which come from the epiblast, and extraembryonic endoderm cells, which are differentiated from the primitive endoderm (hypoblast). Standard ESCs only contributed to yolk sac extraembryonic mesoderm cells (endothelial cells and mesothelial cells)^6^ (Extended Data Fig. 2g). Remarkably, descendants of EPSCs could clearly be found in both the extraembryonic endoderm and the mesoderm cell layers (Extended Data Fig. 2g). These *in vivo* results demonstrated EPSCs’ expanded developmental potential for both the embryo proper and all the major extraembryonic lineages.

To address the possibility of distinct EPSC subpopulations independently contributing to either embryonic or extraembryonic lineages, and to further demonstrate the potency of EPSCs, we tested chimera contribution following the injection of a single EPSC. To this end, each 8C host embryo was injected with a single EPSC (DR10, DR25 and AB2.2) or with a control ESC (AB2.2-2i/LIF) for chimera production. Under the fluorescence microscope, mCherry^+^ cells were found in 7 out of 85 14.5dpc embryos for DR10 EPSCs, 8 out of 77 for DR25 EPSCs, 7 out of 51 for AB2.2-EPSCs, and 10 of 82 for AB2.2-2i/LIF ESCs (Extended Data Fig. 2h). Flow cytometry analysis of the mCherry^+^ chimeras showed EPSC descendants in the embryo proper, and in the extraembryonic tissues in almost every chimera examined (10/11) (Extended Data Fig. 2i). FACS-purified mCherry^+^ placenta cells from EPSC chimeras expressed comparable levels of trophoblast lineage markers (Extended Data Fig. 2j), and contained an 8N polyploid population (Extended Data Fig. 2k). By contrast, injected single AB2.2-ESCs cultured in 2i/LIF contributed poorly to the embryo proper, and little to the placenta in chimeras, as revealed by flow cytometry (Extended Data Fig. 2i).

We formulated EPSCM based on genetic and developmental studies. To dissect the effect of individual inhibitors or their redundancy in EPSCM, we cultured AB2.2 ESCs in various combinations of inhibitors, based on the 2i/LIF medium, for 5 passages before being injected into 8C host embryos. The chimeric blastocysts were co-stained to detect mCherry for the donor cells and Cdx2 for TE contribution. Adding A419259 or XAV939 to the 2i/LIF medium substantially increased TE contributions whereas JNKi and p38i had moderate effects (Extended Data Fig. 2l). The functions of these inhibitors on TE contribution, in particular XAV939 and A419259, were further confirmed by the removal of individual inhibitors from EPSCM (Extended Data Fig. 2l). Although removing CHIR99021 only had a minor effect on the TE contribution (Extended Data Fig. 2l), EPSCs required it for robust proliferation, as revealed in the colony formation assay (Extended Data Fig. 2m). These results thus revealed that the acquisition of TE contribution is possibly an additive effect of modulating more than one kinase or pathway. Importantly, although EPSCM appears to have some redundancy in inhibitor requirement, the percentage of extra-embryonic contribution was highest when all inhibitors were present. On the other hand, the fact that 3-inhibitor combinations, including (2i+A419259, 2i+XAV939, or CHIR99021+A419259+XAV939), were able to confer ESCs with some expanded potential (Extended Data Fig. 2l, 2n) suggests that in the future, EPSCM can potentially be further simplified or optimised. We subsequently examined ESCs cultured in 2i+A419259 medium (2i+A) for their contribution in 14.5dpc chimeras, and detected the descendants of donor cells in the placenta (Extended Data Fig. 2o), which expressed trophoblast markers (Extended Data Fig. 2p), and donor cells in the yolk sac extraembryonic endoderm (Extended Data Fig. 2q).

To understand the molecular characteristics of EPSCs, we profiled the transcriptomes of EPSCs and ESCs in different culture conditions by RNA-seq. In hierarchical clustering, the transcriptomes of the cells segregated by their maintenance conditions, irrespective of their original derivation methods or culture history (Fig. 3a). Using single cell RNA-seq (scRNA-seq), we profiled 84 EPSCs (DR10) to study the molecular heterogeneity of the EPSC culture (Extended Data Fig. 3a). We compared our data with the scRNA-seq dataset of 2i/LIF and M15 ESCs generated on the same platform with an ESC line (G4) (ref. 53). In principal component analysis (PCA), individual cells were segregated by their culture conditions (Extended Data Fig. 3b), demonstrating the global differences between these cells (Extended Data Table 1). Gene ontology term enrichment analysis revealed that 2i/LIF ESCs were enriched in genes of metabolic processes such as oxidative reduction and the electron transport chain (Extended Data Fig. 3c), concordant with a previous report^54^, whereas biological terms related to transcriptional regulation and embryonic development, particularly placental development, were preferentially featured in EPSCs (Extended Data Fig. 3c).

**Figure 3.**
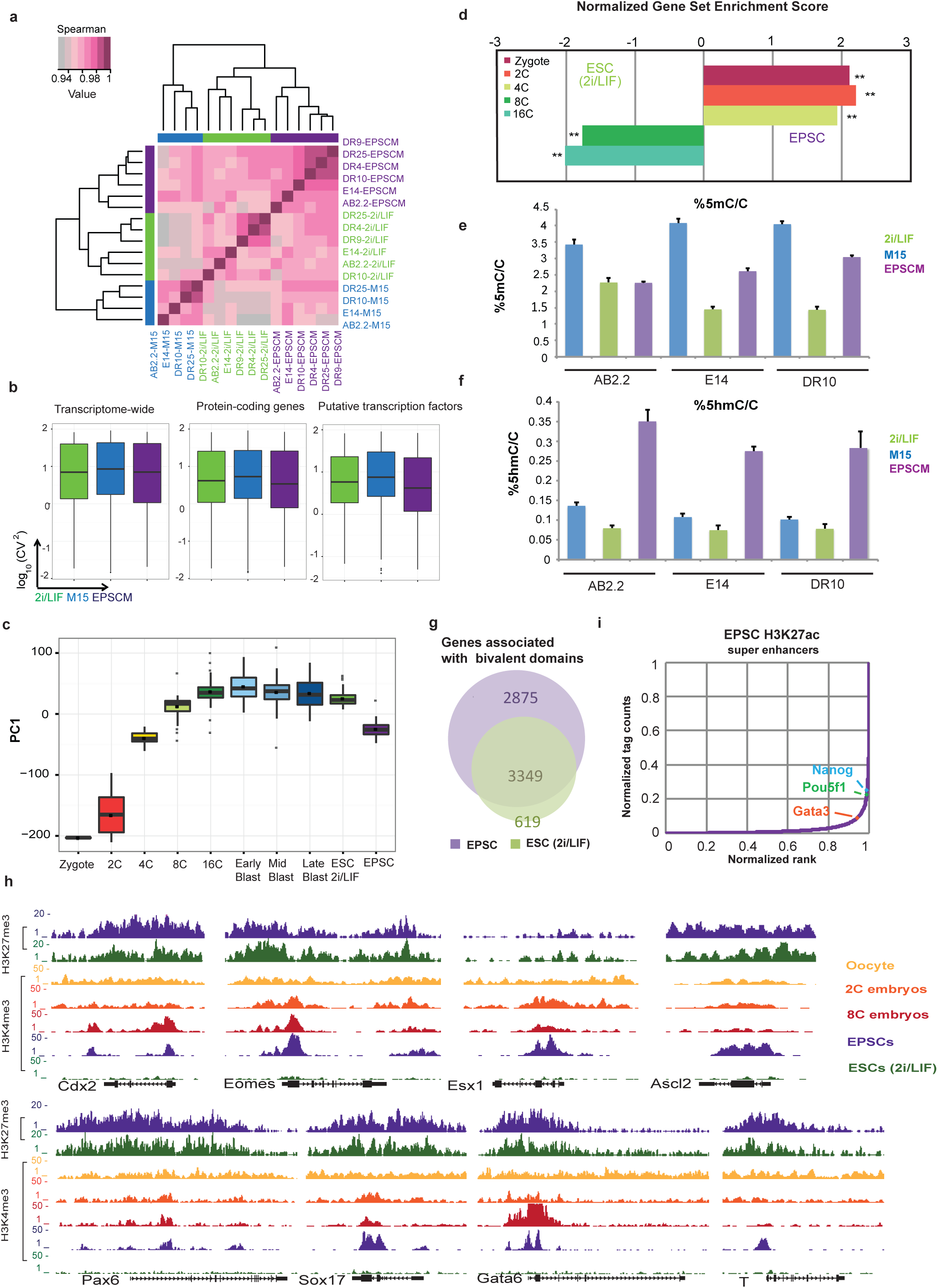
Transcriptomic and epigenomic characteristics of EPSCs. **a.**Unsupervised hierarchical clustering of the population transcriptomes. Standard mouse ESC lines (AB2.2 and E14), ESC lines derived in 2i/LIF media (DR4 and DR9) or EPSC lines derived in EPSCM media (DR25 and DR10) from preimplantation embryos were cultured in M15, 2i/LIF and EPSCM, respectively. The heatmap represents the inter-sample correlation coefficient calculated by the Spearman rank correlation method. Low expression genes with normalized counts less than 5 were excluded. **b.** Boxplots comparing the expression variability of the global transcriptome, protein-coding genes and putative transcription factors in EPSCs and ESCs. The bar in the boxes marks the median of each group. Wilcoxon rank sum test showed significant reduction of squared coefficient of variation of putative transcription factors in EPSCs compared to M15 ESCs (*p* <0.001). **c.** Boxplot comparing the distribution of preimplantation embryos, 2i/LIF ESCs and EPSCs in the first principal component in PCA. The bar and diamond in the boxes mark the median and mean scores of each group. The PCA was performed with differentially expressed genes between EPSCs and 2i/LIF ESCs. **d.** Bar chart showing the normalized enrichment scores of EPSCs and 2i/LIF ESCs in the expression of embryonic stage-specific gene sets. ** nominal *p* value <0.01. The definition of a stage-specific gene set is described in Methods. **e. & f.** Column charts comparing the fractional levels of methylated (**e**) or hydroxymethylated (**f**) cytosine in EPSCs and in ESCs quantified by mass spectrometry. **g.** Venn diagram comparing the number of bivalent histone domains associated genes in EPSCs and 2i/LIF ESCs. **h.** Signal profiles of H3K4me3 and H3K27me3 signals of oocytes, 2C embryos, 8C embryos, EPSCs and 2i/LIF ESCs in selected development genes. Note the similar H3K27me3 profiles in EPSCs and in 2C or 8C embryos. **i.** Individual H3K27ac peak regions were merged and ranked based on their normalized tag counts. Super enhancers were classified as described in Methods. Peak regions associated with *Gata3, Pou5f1* or *Nanog* were marked.

Expression variability of key pluripotency genes (quantified by the coefficient of variation) at the single cell level was compared between EPSCs and 2i/LIF or M15 ESCs, which showed that both EPSCs and 2i/LIF ESCs had similar low variability of these genes (highly homogeneous), unlike the high variability observed in M15 ESCs (Extended Data Fig. 3d). Similarly, EPSCs and 2i/LIF ESCs also had comparable transcriptional variability of protein-coding genes, genes of putative transcription factors as well as global gene expression, which were different from M15 ESCs (Fig. 3b). These scRNA-seq data confirmed that EPSCs had similar transcriptomic homogeneity to 2i/LIF ESCs, and importantly, did not appear to have subpopulations contributing to distinct lineages.

To assess the molecular similarities of EPSCs to *in vivo* preimplantation blastomeres, we retrieved the mouse embryonic development time course single-cell data from Deng et al^55^ for comparison. EPSCs were separated from the native developmental trajectory of blastomeres in the first three principal components (Extended Data Fig. 3e). Yet, the scores of EPSCs were at the range of 4C blastomeres in PC1 (Figure 3c), the component representing the major embryonic development axis. To test whether EPSCs had enriched transcriptomic features of blastomeres compared with standard ESCs, we compiled the top 500 stage-specific genes of each blastomeric stage from Deng et al and compared the expression of these signature gene sets in EPSCs and 2i/LIF ESCs by GSEA. The result showed significantly higher enrichment of early pre-implantation (zygote, 2C and 4C) signatures in EPSCs (Fig. 3d). *Nr5a2* (*Lrh1*) and *Rarg* are specifically highly expressed in 2C-8C blastomeres^55,56^. A recent genome-wide epigenomic profiling study in blastomeres identified them as key determining factors in the segregation of the TE and ICM where *Nr5a2* promotes ICM gene expression^56^. Interestingly, from the scRNA-seq data, EPSCs expressed both genes at high levels whereas only *Nr5a2* was highly expressed in 2i/LIF ESCs (Extended Data Fig. 3d). Similarly, EPSCs, but not 2i/LIF ESCs, had high levels of *Aire, Thap11* (*Ronin*) and *Lin28* (Extended Data Fig. 3d), which are also highly expressed in 2C or cleavage stage blastomeres, respectively^57,58^. Collectively, these results indicate that the transcriptome of EPSCs is enriched with features of early blastomeres. Despite some transcriptomic similarities, it is important to note that EPSCs are *in vitro* cultured cells and are different in many aspects from *in vivo* blastomeres. Finally, the recently reported rare totipotent-like ESC subpopulations (2C-like or MERV-TdTomato^+^, and Hhex-Venus^+^)^36,38^ or *in vivo* reprogrammed iPSCs (iviPSCs)^59^ appear to have trophoblast potential. We showed previously that the 2C-like cells had limited gene expression differences compared to standard ESCs^53^. EPSCs, on the other hand, were distinct from all these cells (Extended Data Fig. 3f). Unlike in 2C-like cells, EPSCs had only limited up-regulation of endogenous retroviral transcripts (Extended Data Fig. 3g).

Blastomeres of 4C-8C embryos have high expression of genes encoding Tet proteins and DNA methyltransferases^55^, and have high levels of 5hmC owing to active demethylation^60^. Further investigation of scRNA-seq data revealed that EPSCs had similar high expression of genes of both methyltransferases (*Dnmt1, Dnmt3a* & *Dnmt3b*) and components involved in active DNA demethylation (*Tet1, Tet2* and *Tdg*) (Extended Data Fig. 3d), whereas 2i/LIF ESCs showed *Prdm14-mediated* down-regulation of *Dnmt3a* and *Dnmt3b* due to ERK signaling inhibition^61^ (Extended Data Fig. 3d and Extended Data Table 1). The expression patterns of the genes encoding DNA methylation/demethylation proteins in EPSCs were reflected on the global cytosine methylation and hydroxymethylation levels. Consistent with previous reports^61,62^, substantially lower levels of DNA methylation were found in 2i/LIF ESCs, compared to that in M15, whereas EPSCs showed an intermediate level (Fig. 3e). Strikingly, much higher levels of hydroxymethyl-cytosine were found in EPSCs (Fig. 3f). The DNA methylome of EPSCs is therefore highly dynamic with active DNA methylation and demethylation.

Besides the dynamic DNA methylome, EPSCs had more genes (6224 vs. 3968) associated with both H3K4me3 and H3K27me3 (bivalent) than 2i/LIF ESCs^54^ (Fig. 3g, Extended Data Fig. 3h, i. and Extended Data Table 2). In EPSCs, bivalency of H3K4me3 and H3K27me3 signals at key pluripotency loci such as *Pou5f1, Sox2* and *Nanog* were highly similar to 2i/LIF ESCs (Extended Data Fig. 3j), in line with the fact that these pluripotency genes were expressed at similar levels in both cell types (Extended Data Fig. 3d). Gene ontology term enrichment analysis showed that EPSC-specific bivalent genes (Extended Data Table 2) were enriched in biological processes of somatic lineage and placental development (Extended Data Fig. 3k), including genetic loci such as *Cdx2, Eomes, Esx1, Ascl2, Pax6, Sox17, Gata6* and *Brachyury* (*T*) (Fig. 3h), providing an epigenetic basis for the observed co-expression of lineage markers in the Hhex^high^ cells in standard ESC culture that show some totipotency features^36^. Remarkably, in line with EPSCs having enriched transcriptomic features of blastomeres, the H3K4me3 patterns of those key developmental loci in EPSCs resembled that of 8C embryos^63^ (Fig. 3h). Long and high intensity H3K27ac domains, i.e. super enhancers, have been associated with master regulators and cellular identity^64^. To further study the epigenetic characteristics of EPSCs, we profiled the genome-wide distribution of H3K27ac in EPSCs (DR10) and identified genes that are associated with super enhancers. As expected, pluripotency master regulators such as *Pou5f1* and *Nanog* were associated with super enhancers in EPSCs (Fig. 3i and Extended Data Fig. 3l), similar to 2i/LIF ESCs. Nevertheless, EPSCs had acquired a unique super enhancer at the *Gata3* locus, an essential regulator for trophectoderm development^65^, which is absent in 2i/LIF ESCs (Fig. 3i and Extended Data Fig. 3l).

Trophoblast stem cell (TSC) lines are derived from the TE or extraembryonic ectoderm of early implantation embryos^66,67^. Since ESCs normally originate from the epiblasts in the ICM, which are already separated from the TE in the blastocyst, it has been challenging to derive stable TSC lines from ESCs. On the other hand, genetic or epigenetic manipulation of mouse ESCs, including transcription factor overexpression or inactivation, or modulation of signal transduction pathways, can make ESCs acquire some potential to differentiate to placental trophoblasts^11,68–70^. However, these TSC-like cells are fundamentally different from TSCs that are derived from mouse embryos or that are reprogrammed from fibroblasts^51,52,70^. In particular, the trophoblast lineage gatekeeper Elf5 is not robustly expressed in these TSC-like cells.

Distinct from standard ESCs, EPSCs possessed enriched transcriptomic features of blastomeres and contributed to the extraembryonic ectoderm and placental trophoblasts in chimeras. Therefore, we endeavoured to establish a stable TSC line from EPSCs, using a direct culture condition switch. We initially cultured AB2.2 EPSCs for 6 days on MEFs in TX medium^71^ containing Fgf4 and TGF-β1, which produced cells of various lineages, including some Cdx2^+^ cell patches morphologically resembling TSC colonies (Extended Data Fig. 4a). To facilitate characterisation of these TSC-like cells from EPSCs, we converted the Cdx2-GFP reporter ESCs^72^ to EPSCs, where a GFP expression cassette is inserted into the *Cdx2* locus to allow tracking Cdx2-expressing cells.

As early as three days after the Cdx2-GFP EPSCs were cultured in TX medium, patches of GFP^+^ cells were visible (Fig. 4a), which were subsequently FACS-purified and re-plated in TX medium. TSC-like colonies from these sorted single cells were picked and expanded to establish stable lines for continuous passaging over 20 times in TX medium (Fig. 4b). Under the same condition, no GFP^+^ cells were detected from 2i/LIF ESCs. The TSC-like cells proliferated similar to the control TSCs, and expressed high levels of TSC genes, including *Cdx2, Elf5, Eomes*, and *Tfap2c*, similar to that in control TSCs (Fig. 4c and Extended Data Fig. 4b). Notably, they did not express the pluripotency gene *Oct4* or the three embryo germ layer markers, *Fgf5, Brachyury* (*T*) or *Gata6* (Fig. 4c). These EPSC-derived TSC-like cells were hence named EPSC-TSCs. Once Fgf4 was withdrawn from TX medium, EPSC-TSCs terminally differentiated into trophoblasts including some polyploid trophoblast giant cells (Extended Data Fig. 4c), and expressed high levels of the differentiated trophoblast markers *Tpbpa, Pl-1, Pl-2, Ctsq* and *Prl2rc2* (ref. 71) (Extended Data Fig. 4d, e). TSCs have the peculiar property to form haemorrhagic tumours when implanted subcutaneously owing to the invasive properties of trophoblast giant cells during implantation^73,74^. EPSC-TSCs formed obvious haemorrhagic lesions in the immunocompromised recipients 7 days after subcutaneous injection (Extended Data Fig. 4f). Histological examination of the lesion sections revealed differentiated trophoblastic giant cells and blood-filled lacunaes, demonstrating an invasive capacity of trophoblasts into host vessels (Fig. 4d and Extended Data Fig. 4g). Finally, immunostaining confirmed Tfap2c^+^ trophoblasts in the sections (Fig. 4d).

**Figure 4.**
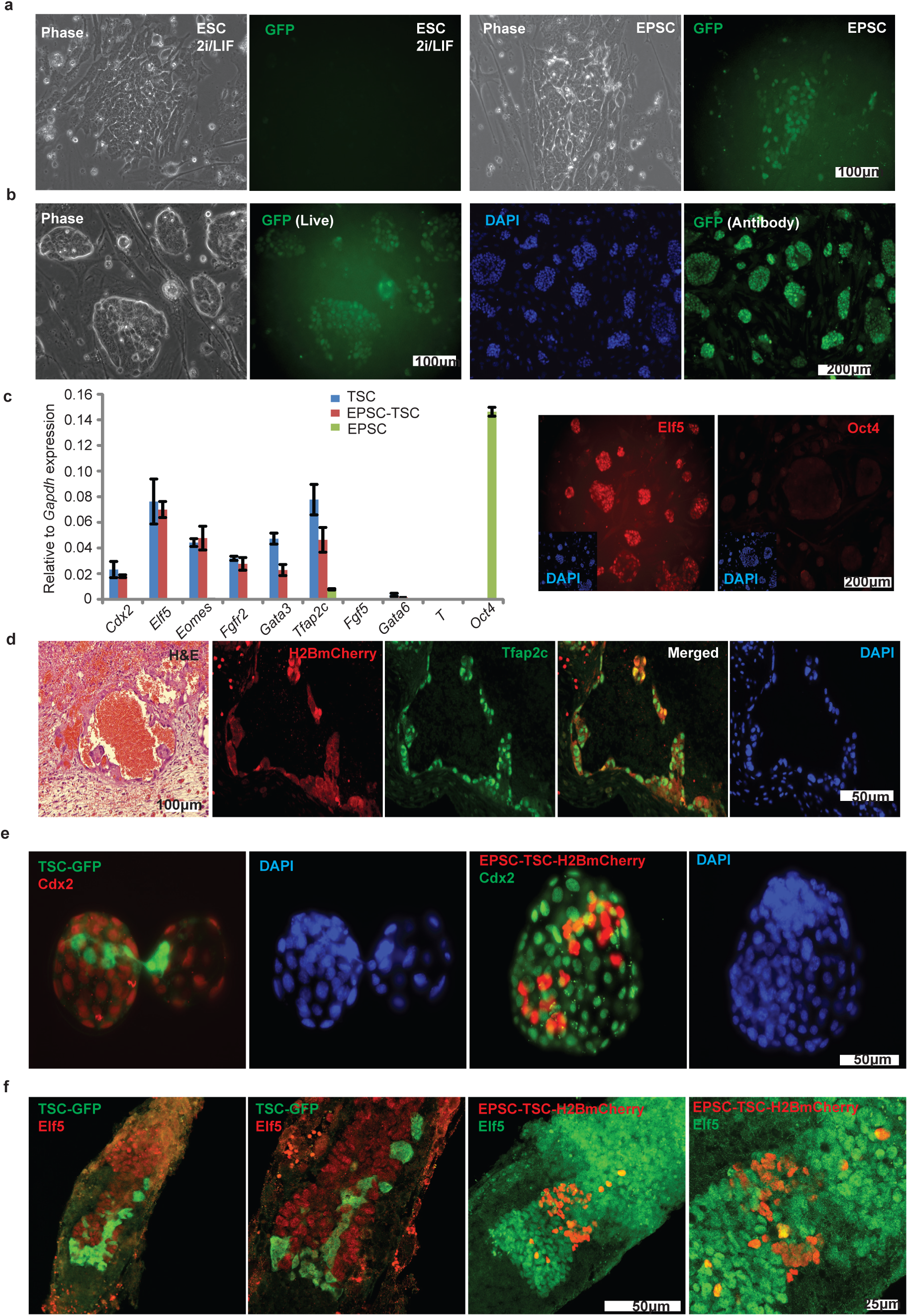
Derivation of TSCs from EPSCs. **a**. Detection of GFP^+^ cells when the Cdx2-GFP EPSCs were cultured in TX medium. No GFP^+^ cells were observed from 2i/LIF ESCs. **b**. Morphology and GFP expression in established Cdx2-GFP EPSC- TSCs. The two images on the left are live images. GFP expression was verified by immunofluroscence staining (images on the right). **c**. qRT-PCR detection of expression of key TSC genes and of *Oct4* in EPSC-TSCs. Expression levels are relative to *Gapdh*. Expression of Elf5 in EPSC-TSCs was also demonstrated by immunostaining. No Oct4 was detected. The insets display DAPI staining. **d**. Sections of hemorrhagic lesions from EPSC-TSCs. More H&E (hematoxylin and eosin) images are in Extended Data Fig. 4g. As expected, EPSC-TSC derived cells stained positively for mCherry. Many mCherry^+^ cells expressed the trophoblast marker Tfap2c. DAPI stains the nucleus. **e**. Localisation of donor cells in the blastocysts from morulas injected with TSCs (cytoplasmic GFP) or EPSC-TSCs (H2B-mCherry). DAPI stains the nucleus. Some donor cells expressed Cdx2. **f**. Contribution of TSCs (cytoplasmic GFP) or EPSC-TSCs (H2B-mCherry) in the extraembryonic ectoderm region marked by Elf5 in immunostaining. DAPI stains the nucleus. The stained chimera embryos were examined using confocal microscopy. All experiments were repeated at least three times.

To determine whether EPSC-TSCs can function properly in their native environment, we injected the H2B-mCherry-expressing EPSC-TSCs into recipient 8C embryos, which were further cultured for 48 hours prior to staining for mCherry and Cdx2 expression to facilitate localization of the injected cells. Similar to TSCs (expressing a cytoplasmic GFP from a constitutive promoter)^66^, the mCherry^+^ EPSC-TSCs were found in the TE or the outer layer of the blastocysts (TSCs: 31 out of 40 blastocysts; EPSC-TSCs: 127 out of 163 blastocysts) with some cells expressing Cdx2 (Fig. 4e). We implanted the injected embryos into recipient mice, and collected E6.5 embryos for analysis. Both the control TSCs (GFP^+^) and the mCherry^+^ EPSC-TSCs contributed to the Elf5^+^ extraembryonic ectoderm region (chimeras and total embryos: TSCs, 19 out of 45; EPSC-TSCs, 33 out of 94) (Fig. 4f). These results proved that *bona fide* TSCs were directly derived from EPSCs by a simple culture condition switch, and further validated the expanded potential of EPSCs.

These results thus provide clear proof that deriving new stem cells with expanded potential can be achieved. It is envisaged that the successful establishment of mouse EPSCs from cleavage stage embryos and by conversion from ESCs/iPSCs will offer a unique opportunity for the study of the earliest stages of embryo development. Furthermore, the insights gained herein may accelerate the establishment of cultures of similar stem cells from other mammalian species for which embryonic stem cells or iPSCs are still not available.

## Acknowledgments

We thank colleagues of RSF (Brendan Doe, Stuart Newman and Evelyn Grau and others), Yvette Hooks, Sequencing (Natalie Smerdon) and FACS core facilities (Bee Ling Ng and Jennifer Graham) at the Sanger Institute, the animal facility at CRUK-CI for excellent technical support, Dr. Myriam Hemberger for providing TSCs and Dr. Sebastian Gerety for the fluorescence stereo microscope. We thank Dr. Jong Kyoung Kim for informatics advice, Stephen Rice for helps on DNA bisulfite sequencing analysis and Dr. David Goulding for confocal imaging facility. We thank Dr. Jennifer Nichols for their comments and for female NOD ESCs and TSCs. We thank Professor Jamie Thomson for comments. We appreciate critical comments from Professors Jamie Thomson and Elizabeth Robertson. D.R. is a recipient of the Wellcome Trust Clinical PhD Fellowship for Academic Clinicians. L.A. is a recipient of a Ph.D. fellowship from the Portuguese Foundation for Sciences and Technology, FCT (SFRH/BD/84964/2012). Y. T. was supported by a Japan Society for the Promotion of Science fellowship. A.C.W. was supported by the National Institute of Health Research (RP-PG-0310-10002). M. E-M is supported by an EMBO Fellowship (ALTF938-2014) and Marie Sklodowska-Curie Individual Fellowship. W. R. acknowledges funding from BBSRC (BB/K010867/1) and Wellcome Trust (095645/Z/11/Z). L. Lu is supported by the National Natural Science Foundation of China (31370904 and 30972691). P. L. thanks Professors Mike Stratton, Allan Bradley, Neal Copeland, Nancy Jenkins and James Lupski for their encouragement in these experiments. P. L. lab is supported by the Wellcome Trust (grant number 098051).

## Author contributions

D. J. R. and W. W. developed EPSCM, derived mouse EPSC lines, performed the *in vitro* and *in vivo* differentiation assays, edited the manuscript and made figures. J. Y. produced mouse iPSCs, performed Western blots, immunostaining of 5.5-7.5dpc embryos, placenta sections, yolk sac, and sorted placenta cells, analysed preimplantation embryos in culture and immunostaining, investigated minimal requirement of inhibitors, deriving and characterising EPSC-TSCs, made the final figures and edited the manuscript. J. C. T. carried out the single-cell RNA-seq experiment, analyzed the RNA-seq and ChIP-seq data, and produced the genomics figures and wrote the manuscript. X. G., L. A., Y. Y., J. W. and C. W. performed experiments. L. C. interpreted the placental and teratoma slides. F. Y. and B. F. karyotyped the EPSC and ESC lines. G. L. performed most of the EPSC and ESC injections. H. M. performed independent mouse EPSC conversion and injection experiments at Nakauchi lab. M.W. assisted embryo production. Z. Z. performed confocal imaging and interpretation. J. B., R. R. and X. Z. provided microinjection resources. A. K. and S. T. provided single-cell RNA-seq data from standard ESCs. A.W. performed EPSC differentiation, Y.T. carried yolk sac immunostaining and embryo dissection, and B. G. provided supports for A. W and Y. T. M. E-M. and W. R. provided whole genome DNA methylation data. L. L. contributed intellectually and experimentally. P. L. conceived the concept, designed the studies, wrote the manuscript, and supervised the overall research project and manuscript preparation.

## Extended Data Figures

**Extended Data Figure 1 | Derivation and functional characterization of EPSC lines from pre-implantation embryos, by converting standard ESCs and by reprogramming somatic cells. a.** Phase images showing development of 8C mouse embryos in EPSCM or in M15. Note that in the EPSCM embryos, the original blastocoel cavity is now filled with cells and that the embryos hatch much later than embryos in M15 medium. **b**. More phase images of 8C embryo development in EPSCM on SNL feeders or feeder-free. **c**. Normal development of 4C embryo in M15. Oct4 expression is detected by immunostaining in 4C blastomeres whereas Cdx2 is expressed in the 8C embryo. In the blastocyst, Oct4 is primarily in the ICM and Cdx2 is restricted to the TE. DAPI stains the nucleus. **d**. Detection of Ki67 for proliferative cells in 8C embryos cultured in EPSCM. Note the complete absence of nuclear Oct4 staining in the mid “filled” blastocyst. **e**. Detection of cleaved Caspase 3 for apoptotic cells in 8C embryos cultured in EPSCM. Note the complete absence of Oct4 staining in the mid “filled” blastocyst. Oct4^+^ cells re-appear in the late “filled” blastocyst. **f**. EPSCs derived from preimplantation embryos express pluripotency markers. Analysis of pluripotency markers in two preimplantation embryo-derived EPSC lines (DR25 and DR10), one somatically reprogrammed iPSC-EPSC line and two preimplantation embryo-derived 2i/LIF ESC lines (DR4 and DR9). Relative expression of pluripotency genes was normalized to *Gapdh* in the qRT-PCR assay. Data are the mean ± s.d. from three experiments. Relative expression of the lineage-specific genes is normalized to *Gapdh*. Data are the mean ± s.d. from three experiments. **g**. Expression of lineage-specific genes in the cells described in **f**. Relative expression of these genes is normalized to *Gapdh*. Data are mean ± s.d. Experiments were repeated three times. **h**. Spectral karyotyping of DR10-EPSCs (normal karyotype in 8 out of 10 metaphases). **i**. Mature teratomas from DR10-EPSCs. Left panel: neural tube-like structures from the ectoderm (neuroectoderm); middle panel: cartilage from the mesoderm; right panel: a gland of possible gastrointestinal type arising from the endodermal layer and muscle fibers derived from the mesoderm. The sections were stained with hematoxylin-eosin. **j**. A chimera derived from DR10-EPSCs. **k.** The mCherry^+^ EPSCs show extensive contribution to the 14.5dpc gonad, similar to 2i/LIF ESCs. **l**. Additional images of EPSC contribution in the TE of the blastocyst. The two images on the left are merged live images (phase and mCherry) the contribution of mCherry (cytoplasmic)-labeled 2i/LIF ESCs or DR10-EPSCs. The three images on the right are immunostaining ones to detect H2B-mCherry-expressing AB2.2 ESC-EPSCs. The image on the far right shows co-expression of H2B-mCherry and Cdx2 in TE cells. **m**. Rex1-GFP reporter ESC colonies in M15, 2i/LIF or EPSCM on SNL feeders. **n**. Expression of the reporter GFP in ESCs cultured in EPSCM (passage 3). Left panel, *Oct4-GFP* ESCs (E14tg2a) cultured in M15, 2i/LIF or EPSCM. In all conditions, *Oct4-GFP* expression is comparably detected in flow cytometry. Right panel: *Rex1*-GFP ESCs (AB2.2 background) also have comparable levels of *Rex1* expression (GFP) when cultured in EPSCM or when subsequently re-cultured in 2i/LIF. Negative control: wild-type ESCs. **o**. Expression of pluripotency genes or lineage-specific genes in ESCs cultured in EPSCM (ESC-EPSCs). Relative expression of these genes is normalized to *Gapdh*. Data are mean ± s.d. Experiments were repeated three times. **p**. Diagram of *Oct4* distal and proximal enhancer luciferase reporter constructs. DE: the *Oct4* distal enhancer; PE: the *Oct4* proximal enhancer; MiniP: the minimum promoter. The *Oct4* distal enhancer is active in ESCs cultured in EPSCM. Luciferase activities of the *Oct4* DE and PE constructs were normalized to that of the MiniP construct in the same cell type. Data are mean ± s.d. from three experiments. **q**. Differentiation of ESC-EPSCs *in vitro*. These cells can differentiate into all three germ layers *in vitro*. β3-Tubulin, α-Smooth muscle actin (SMA) and Gata4 antibodies were used to detect the differentiated cell types representative of the three germ layers. **r**. A male chimeras from ESC-EPSCs, and germline transmission of the *Rex1-GFP* allele from the chimera. **s**. Reactivation of X chromosome in female ESC-EPSCs. EPSCs and the cells differentiated from EPSCs were co-immunostained for H3K27me3 and Oct4. No discrete H3K27me3 foci were found in EPSCs. Once the cells became differentiated, the foci appeared in almost all cells. **t**. Effects of small-molecule inhibitors on their respective targets. p-ERK: phosphorylated ERK; p-SRK: phosphorylated SRK; p-P38: phosphorylated P38; p-JNK: phosphorylated JNK. α-tubulin was used as the loading control. Note the substantial increase of AXIN1. EPSCM has 20% of KSR so KSR-2i/LIF serves as the proper control. **u**. XAV939 in EPSCM stabilizes AXIN1 by inhibiting its ubiquitination. EPSCs had considerably elevated levels of AXIN1 compared with controls as shown above, which caused accumulation of phosphorylated β-catenin in both the cytoplasm and nuclei and decreased active β-catenin in the nucleus. α-tubulin and Histone H3 were used as the loading control of cytoplasmic and nuclear proteins, respectively. **v**. TopFlash luciferase assay confirmed the markedly reduced β-catenin-LEF/TCF activity in EPSCs. **w**. Western analysis shows increased p-STAT3 in ESC-EPSCs with increased concentrations of LIF. The chart on the right describes up-regulation of LIF pathway downstream genes in ESC-EPSCs in response to LIF stimulation. **x**. Signalling pathway dependence in ESC-EPSCs. One hundred ESCs cultured in M15 or 2i/LIF, or EPSCs were plated in their respective medium, and cultured with inhibitors of JAK (JAK inhibitor 1), FGFR (SU5402), TGFBR (A83-01) or ALK5 (SB505124). Alkaline phosphatase-positive (AP^+^) colonies were scored after 10 days. The Jak inhibitor substantially reduced the number of AP^+^ colonies from these cells. By contrast, ESCs cultured in EPSCM did not appear to be sensitive to FGFR or TGFBR inhibitors, similar to 2i/LIF ESCs. Data are the mean ± s.d. from four independent repeats. *: p<0.05.

**Extended Data Figure 2 |Contribution of EPSCs in chimeras. a.** Whole mount immunostaining of 6.5-7.5dpc chimeras for H2B-mCherry. Descendants of EPSCs were found in both the embryo proper and extra-embryonic ectoderm regions. The embryos were labelled individually as 1’ (2i/LIF ES cells), and 2’, 3’ or 4’ (EPSC). The three confocal images in the bottom panel are #2 embryo in Figure 2a, the circled areas of embryos 2’ and 3’. **b**. Whole-mount fluorescence imaging of 14.5dpc chimeras from morula embryos injected with ESCs or EPSCs. Negative control: wild-type embryo. **c.** Detection of polyploid placenta cells by flow cytometry. A distinct population of 8N cells is present in both mCherry^+^ and mCherry^−^ placenta cells from an AB2.2-EPSC chimera. Fetal brain cells were used as the negative control. No mCherry^+^ 8N cells could be detected in the placenta from 2i/LIF ESCs due to the low number of mCherry^+^ cells. **d.** Placenta sections of 14.5dpc chimeras from either ESCs or EPSCs (AB2.2, mCherry^+^) were co-immunostained for mCherry and Tfap2c. mCherry^+^ cells were detected in the EPSC chimera placenta sections, and some of them were positively stained for both mCherry and Tfap2c. The area labeled in DAPI indicates the position of the images in the placenta sections. **e**. mCherry^+^ cells were sorted from 14.5dpc EPSC chimera placenta, cytospun to polylysine coated slides, and co-immunostained for mCherry and trophoblast markers GCM1, Ezrin or CK7 (Cytokeratin 7). The arrows point to mCherry^+^ cells that are positively stained for these markers. **f**. mCherry^+^ EPSCs were injected into *ROSA26-GFP-SB10* blastocysts. The chimeras were collected at 14.5dpc. Both mCherry^+^ and mCherry^−^ placenta cells were sorted, genomic DNA was extracted for PCR. *SB10* DNA (*Sleeping Beauty transposase* gene) should be amplified from the host cells, and mCherry should be amplified from the donor cells. The mCherry^+^ placenta cells had robust *mCherry* DNA amplification, but did not have any detectable host cell DNA, as no SB10 DNA amplification was found. In contrast, weak mCherry signal was found in mCherry^−^ cell sample. This is likely due to the silencing of the CAG promoter in placenta cells derived from the mCherry^+^ donor cells. Amplification of a region in the *Oct4* distal enhancer region serves as the genomic DNA quality and PCR control. **g.** Yolk sac sections of 14.5dpc chimeras of either ESCs (Panels 1-3) or EPSCs (Panels 1’-3’). Due to initial high autofluorescence of FITC-conjugated antibody for mCherry in the yolk sac, we analysed yolk sac sections of chimeras from injecting Rosa26-GFP reporter EPSCs or ESCs. The original GFP fluorescence signal was quenched due to fixation. A CF660C conjugated antibody against GFP was used to detect donor cells. DAPI staining for the nucleus. GFP^+^ cells were found in extraembryonic mesoderm cells (endothelial and mesothelial) in the yolk sac of chimeras of both EPSCs and ESCs. In contrast, GFP^+^ cells were also found in the extraembryonic endoderm layer (extraembryonic-visceral-endoderm-derived) in the yolk sac of EPSC chimeras. **h-k**. Chimers of injecting a single EPSC. **h**. Whole-mount fluorescence imaging of representative 14.5dpc chimeras from 8C embryos injected with a single EPSC or ESC. Negative control: wild-type embryo. **i**. Flow cytometry analysis of mCherry^+^ placenta cells. **j**. Expression of trophoblast genes in sorted mCherry^+^ and mCherry^−^ placenta cells from an EPSC chimera of a single EPSC. Expression was normalized to fetal brain expression. Data are the mean ± s.d. Sorting was performed as in Fig. 2b. **k**. Detection of polyploid placenta cells. A distinct population of 8N cells was found in both mCherry^+^ and mCherry^−^ placenta cells in an EPSC chimera. **l**. Effects of individual inhibitors on TE contribution. Mouse ESCs (mCherry^+^ AB2.2) were cultured in various combinations of inhibitors for at least 5 passages before injection into 8C embryos for blastocyst development. Blastocysts were scored for donor cell TE contribution. A: A419259; X: XAV939; JNKi: JNK Inhibitor VIII; p38i: SB203580; CH: CHIR99021. All culture conditions contained LIF. For each culture condition, 48-50 embryos were stained and scored. **m**. The effect of CHIR99021 (CH) in EPSCM on cells. One thousand EPSCs cultured in either EPSCM or EPSCM minus CH were plated for single cell colony formation. AP^+^ colonies were scored on day 7. **n**. ESCs cultured in minimum sets of inhibitors express pluripotency markers and Low levels of lineage-specific genes. Relative expression of pluripotency genes was normalized to *Gapdh* in the qRT-PCR assay. Data are the mean ± s.d. from three experiments. **o**. Flow cytometry analysis of mCherry^+^ placenta cells from 14.5dpc chimeras of ESCs cultured in 2i+A medium. To minimize residual background cell sorting, we also used a GFP channel to exclude autofluorescence. **p**. Expression of trophoblast-enriched genes in cells sorted from the placenta of chimeras. The mCherry^+^ placenta cells are derived from ESCs cultured in 2i+A medium as in **o**. Expression was normalized to *Gapdh*. Data are the mean ± s.d. **q**. Yolk sac sections of 14.5dpc chimera of EPSCs (2i+A) or the negative control. Immunostaining and imaging were performed in the same way as in Extended Data Fig. 2g. The original mCherry fluorescence signal was quenched due to fixation. Donor cells were stained by a CF660C conjugated antibody against mCherry (Panels 1 and 1’). DAPI staining was used to detect the nucleus (Panels 2 and 2’). Donor cell-derivatives were found in both the extraembryonic mesoderm (endothelial and mesothelial cells) and extraembryonic endoderm cells in the yolk sac of EPSC chimera, but not in the yolk sac of the negative control chimera (3 and 3’).

**Extended Data Figure 3 | Single cell RNA-seq analyses and epigenomic characteristics of EPSCs. a.** Scatterplot showing the relationship of EPSC gene expression variability with expression levels in scRNA-seq data. The magenta dots represent genes that showed significantly higher variability (adjusted *p* <0.1) than would be expected from the external RNA spike-ins (blue dots). **b.** Three-dimensional scatterplot showing the separation of 2i/LIF ESCs, M15 ESCs and EPSCs in PCA. Each dot represents a cell. **c.** Bar chart showing gene ontology terms enriched in differentially expressed genes between 2i/LIF ESCs and EPSCs in scRNA-seq data. **d.** Violin plots comparing expression of selected pluripotency regulators, DNA de/methylation regulators and 2-cell embryo-associated genes in 2i/LIF ESCs, M15 ESCs and EPSCs in scRNA-seq data. The dot represents the mean expression level. **e.** Three-dimensional scatterplot showing the separation of different stages of preimplantation embryonic blastomeres, 2i/LIF ESCs and EPSCs in PCA of scRNA-seq data. **f.** Three-dimensional scatterplot showing the separation of Hex-Venus+ 2i/LIF ESCs, MERV-TdT^+^ ESCs, in vivo iPSCs, 2i/LIF ESCs, M15 ESCs and EPSCs in PCA of bulk RNA-seq data. **g.** Column chart comparing the relative expression levels of MERV transcripts in EPSCs and MERV-TdT^+^ ESCs to MERV-TdT^+^ ESCs. **h.** Signal intensity distribution of EPSC H3K4me3 and H3K27me3 modification over gene bodies and 3kb up/downstream of transcription start/end sites. Genes are classified into “Very High”, “High”, “Low” and “Very Low” depending on their length-corrected mean-normalized count quartiles in the single-cell RNA-seq dataset. **i.** Distribution of H3K27me3 (left) and H3K4me3 (right) signals at gene promoters (+/−3kb from TSS). The promoters were ranked based on their length-corrected mean-normalized count levels in the single-cell RNA-seq dataset. The signals were quantified as read count per million mapped reads. **j.** Profiles of H3K4me3 and H3K27me3 signals of 2i/LIF ESCs and EPSCs in selected pluripotency-associated genes. **k.** Bar chart showing selected biological gene ontology terms enriched in EPSC-specific bivalent genes. Only terms relevant to development are shown here. **l.** Profiles of H3K27ac signals of oocyte, 2C embryos, 8C embryos, 2i/LIF ESCs and EPSCs in selected super enhancer-associated genes, including *Pou4f1, Klf5* and *Gata3* loci. Note that the super-enhancer at the *Gata3* locus is present in EPSCs and in 8C embryos but not in 2i/LIF ESCs.

**Extended Data Figure 4 | Derivation and characterization of EPSC-TSCs. a.** Immunofluorescence staining for GFP in EPSCs or 2i/LIF ESCs cultured in TX medium for 6 days. Small patches of Cdx2^+^ cells were only detected in differentiated EPSCs. DAPI stains the nucleus. The Cdx2^+^ cells account for around 1.0% of total cells in flow cytometry. **b.** Expression of Cdx2, Eomes and Tfap2c in established EPSC-TSCs detected by immunostaining. Four EPSC lines were established from Cdx2-GFP EPSCs. **c**, Phase images of EPSC-TSCs in differentiation medium (RPMI 1640 plus 20% serum without FGF4 and heparin) for the indicated days. Arrows indicated possible polyploid trophoblasts. **d.** qRT-PCR analysis of differentiated EPSC-TSCs or the control TSCs at indicated days. Cdx2 was down-regulated while expression of mature trophoblast genes was up-regulated. Expression levels are relative to *Gapdh*. **e**. Immunofluorescence staining of differentiated EPSC-TSCs or TSCs for placenta lactogen-1 (PL-1) (day 8). The insets display DAPI staining. **f.** Representative images of hemorrhagic lesions in NSG mice 7 days after subcutaneous injection of EPSC-TSCs or TSCs. **g**. H&E sections of hemorrhagic lesions of EPSC-TSCs or TSCs. The images show that the well defined haemorrhagic lesions are lined by large and occasionally multinucleated pleomorphic cells with abundant cytoplasm. These large cells outline spaces filled with blood and show the features of trophoblast giant cells.

